# Experimental evidence for a microbial origin of reduction spots in red beds

**DOI:** 10.64898/2026.01.12.699011

**Authors:** Sev Zielinska, Naomi Felton, Philip Vixseboxse, Sean McMahon

## Abstract

Perseverance rover recently discovered sedimentary rocks reddened by ferric oxides and peppered with bleached spots lacking these oxides. Some of these spots are associated with phosphate, iron sulfide minerals, and organic matter, and are regarded as “potential biosignatures”, suggestive of microbial iron-and sulfate-reduction and organic matter oxidation. Similar mm–cm-scale “reduction spots” occur in many ancient “red beds” on Earth. Although terrestrial reduction spots are widely considered biogenic, the available evidence is not decisive, and the proposed microbial mechanism of spot formation has not been tested experimentally. Here, we report a successful laboratory demonstration of bleached spot-formation in ferruginous sediment. Mm–cm-scale rounded bleached spots appeared within weeks on the underside of anaerobic sand–ferrihydrite slurries inoculated with mixed microbial communities obtained from the reducing zones of Winogradsky columns. The spatial and temporal distribution of observed bleaching events, which did not occur in sterile controls, are best explained by a microbially induced process, and DNA sequencing confirms that bacteria of iron-reducing genera are abundant in the bleached areas. These results strongly support the longstanding hypothesis that microbial colonies can indeed generate visibly bleached reduction spots in ferruginous sediments and rocks. Further experiments are needed to establish whether and how non-biological processes can mimic these features, and to search for features that disambiguate biogenic and abiogenic reduction spots.

## Introduction

In mid-2024, during its exploration of the Neretva Vallis channel leading into Jezero Crater, Mars, NASA’s *Perseverance* rover encountered bleached (ferric-oxide-depleted) spots in the ferric-oxide-rich sedimentary rocks of the Bright Angel formation (Hurowitz et al., 2025). Similar “reduction spots” are common in ferric-oxide-rich “red beds” on Earth and have long been considered a potential biosignature (e.g., Hofmann, 1990; Spinks et al., 2010).

*Perseverance* found two populations of spots. Brick-red mudstones of the Mount Spoonhead member contained pale greenish mm-scale halos around mottled dark cores. Pink mudstones of the Cheyava Falls member contained organic matter and pale mm-scale “leopard spots” depleted in ferric iron, enriched in ferrous iron, and enriched in sulphide relative to the matrix. These spots were lined with dark ferrous phosphate, also present in accompanying μm-scale “poppy seeds”. Both populations of spots are best explained by localized reductive dissolution and loss of ferric iron; at Cheyava Falls, reduction of iron and sulphate may have been coupled to the oxidation of organic matter, liberating phosphate (Hurowitz et al., 2025). *Curiosity* has also observed features resembling reduction spots, with white halos and dark cores, in the slightly pink Carolyn Shoemaker formation of Gale Crater (Seeger and Grotzinger, 2024).

On Earth, reduction spots are thought to form when microorganisms couple the oxidation of organic matter to the reduction of iron in rocks and sediments, creating a redox front that expands radially (e.g., Hofmann, 1990). However, an abiotic origin can rarely be excluded, and evidence for biogenicity is typically indirect. For example, Spinks et al. (2010) reported sulphide δ^34^S values in a population of Triassic reduction spots significantly lighter than co-eval sulfates, suggestive of microbial sulfate reduction but within the range of (thermochemical) abiotic sulfate reduction. Similarly, in a global sample of Mesoproterozoic–Recent reduction spots, McMahon et al. (2018) found a pattern of progressive enrichment in δ^238^U from matrix to halo to core, consistent with the enzymatic fractionation of uranium isotopes by metal reducing bacteria (Stylo et al., 2015) but not unequivocally so (Brown et al., 2018).

Because the redox reactions that occurred at Cheyava Falls could have been driven either by microorganisms or by abiotic chemistry, Hurowitz et al. (2025) describe them as “potential biosignatures”. A phenomenon is a “potential biosignature” if, firstly, “biological processes are a known possible explanation” and, secondly, “potential abiotic causes have not yet been reasonably explored and ruled out” (Gillen et al., 2023). Arguably, reduction spots have not hitherto met the first criterion (let alone the second) because it has never been demonstrated that biological processes can produce them. Here, we report experimental evidence that microbes can indeed produce visibly bleached reduction spots in ferruginous sediment.

## Methods

### Microbial inoculum

Sediment was collected in polypropylene cylinders from the Hermitage of Braid and Blackford Hill Local Nature Reserve on 12 May 2025. Soil was collected at 55°55′31.5”N 3°11′39.5”W at ∼5 cm depth; dark pond mud from the bank of Blackford Pond at 55°55′32.9”N 3°11′39.6”W. Sediment was sieved through a 1-mm mesh.

Six Winogradsky columns were prepared in polypropylene cylinders (8.5 cm diameter, 18 cm height). Each contained (1) an enriched slurry of 1.67 g CaCO_3_, 1.67 g CaSO_4_, 6.67 g blended newspaper and tap-water (1:3), and 33.3 mL of the environmental sample, following Lalla et al. (2021); and (2) 350g of sample sediment and 100 mL of tap-water. Four columns also contained 0.5 g ferrihydrite (synthesized according to Smith et al., 2012). The columns were capped with clingfilm and rubber bands, and labelled according to sample source (S, soil; P, pond), presence (F) or absence (NF) of ferrihydrite, and growth conditions (L, light, D, dark), i.e., SNFL, SFL, SFD, PNFL, PFL, PFD. “Light” columns were placed on a south-facing windowsill and exposed to a natural day/night cycle.

Deep black layers typical of microbial sulfate reduction (Rogan et al., 2005) developed at the base of all six columns by day 20 and were respectively 4.6 cm (PFD), 5.9 cm (PFL), 6.3 cm (SFD, SFL), 9.3 cm (PNFL), and 9.2 cm (SNFL) thick (**Supplementary Figure 1**).

### Sediment bleaching experiments

In Experiment 1, 57 days after the columns were established, 1 mL of inoculum was transferred by sterile Pasteur pipette from the upper region of each black layer in each column into an autoclaved 1-mm ball of absorbent cotton wool (see **Figure 1** and below). One cotton wool ball received 1 mL of sterile pond water instead of a microbial inoculum. In Experiment 2 (day 114), the black layer in columns SFL and PNFL were sampled as before. In addition, a sterile control was prepared whereby the inoculum from the column was passed through a 0.22 µm nylon syringe filter before soaking into the cotton ball. In Experiment 3 (day 137), the black layers in columns SFL and PNFL were again sampled as before. An additional experiment was performed using a pure culture of the iron-reducing bacterium *Geobacter bemidjiensis* (DSMZ) cultivated on Geobacter Medium 579.

**Figure 1.**
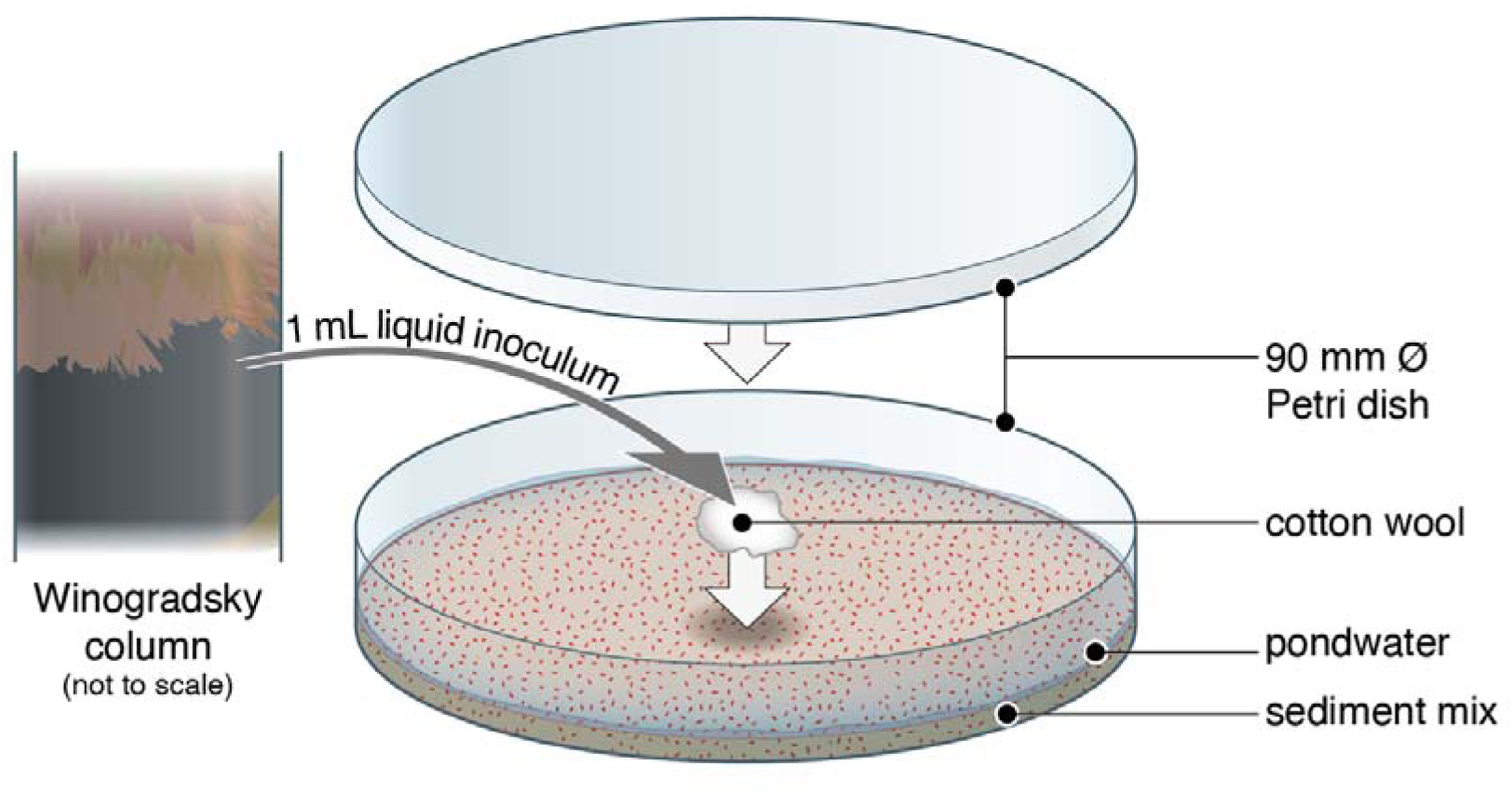
Schematic illustration of experimental set-up. The sediment mix consisted of 98 wt% quartz powder and 2 wt% synthetic ferrihydrite. Dishes were sealed with parafilm and placed in an anaerobic chamber under 20% CO_2_, 80% N_2_. See Supplementary Figure 1 for photographs of Winogradsky columns.

Under a laminar flow hood, “quartz sand” powder (particle size <63 µm, Sigma-Aldrich; sterilised at 150°C for 16 hr in a Teflon bottle) was mixed with 2 wt% ferrihydrite following Vixseboxse et al. (2024) and dispensed into 90-mm-diameter Petri dishes (22 g per dish; Experiments 1 and 2) or 40-mm dishes (3 g per dish; Experiment 3). Each dish was moistened with 10 mL of sterilized (filtered < 0.22 µm) diluted pond water (collected from Blackford Pond, 11 June 2025) using a sterilized spray-bottle, producing a <1 mm supernatant above the saturated sediment. Dishes exchanged gas for three days in an anaerobic chamber (Coy; Vinyl Anaerobic Airlock Chamber, Michigan, USA; 20% CO_2_, 80% N_2_ atmosphere).

The inoculated cotton balls were placed gently on top of the sediment in the middle of each dish using sterile tweezers. All dishes were parafilm-sealed. Water lost by evaporation was replenished twice monthly to maintain the supernatant. Dishes were infrequently removed from the anaerobic chamber during the experiment for photography (∼3 minutes) and flat-bed scanning (∼5 minutes, using a XEROX AltaLink C8035 scanner).

The methods used for DNA extraction and sequencing, resin impregnation, and electron microscopy (including energy-dispersive X-ray spectroscopy, EDX) are described in **Supplementary Information**.

## Results

### Development of bleached spots

In Experiment 1, bleaching occurred only in dishes inoculated from columns SFL and PNFL (**Table 1**). On day 22, bleached spots were noticed on the undersides of the sediment in both dishes, corresponding with the positions of the cotton wool (**Figure 2**). In PNFL, the spot was circular, faint with indistinct margins, and ∼6 mm across. By day 36, it had expanded to ∼8 mm, become less faint, and acquired a slight orange halo; no further change occurred (**Figure 2**). Patches of rust-orange material formed at the sediment-water interface in (only) the same two dishes, directly above the bleached zones (**Supplementary Figure 2**).

**Figure 2:**
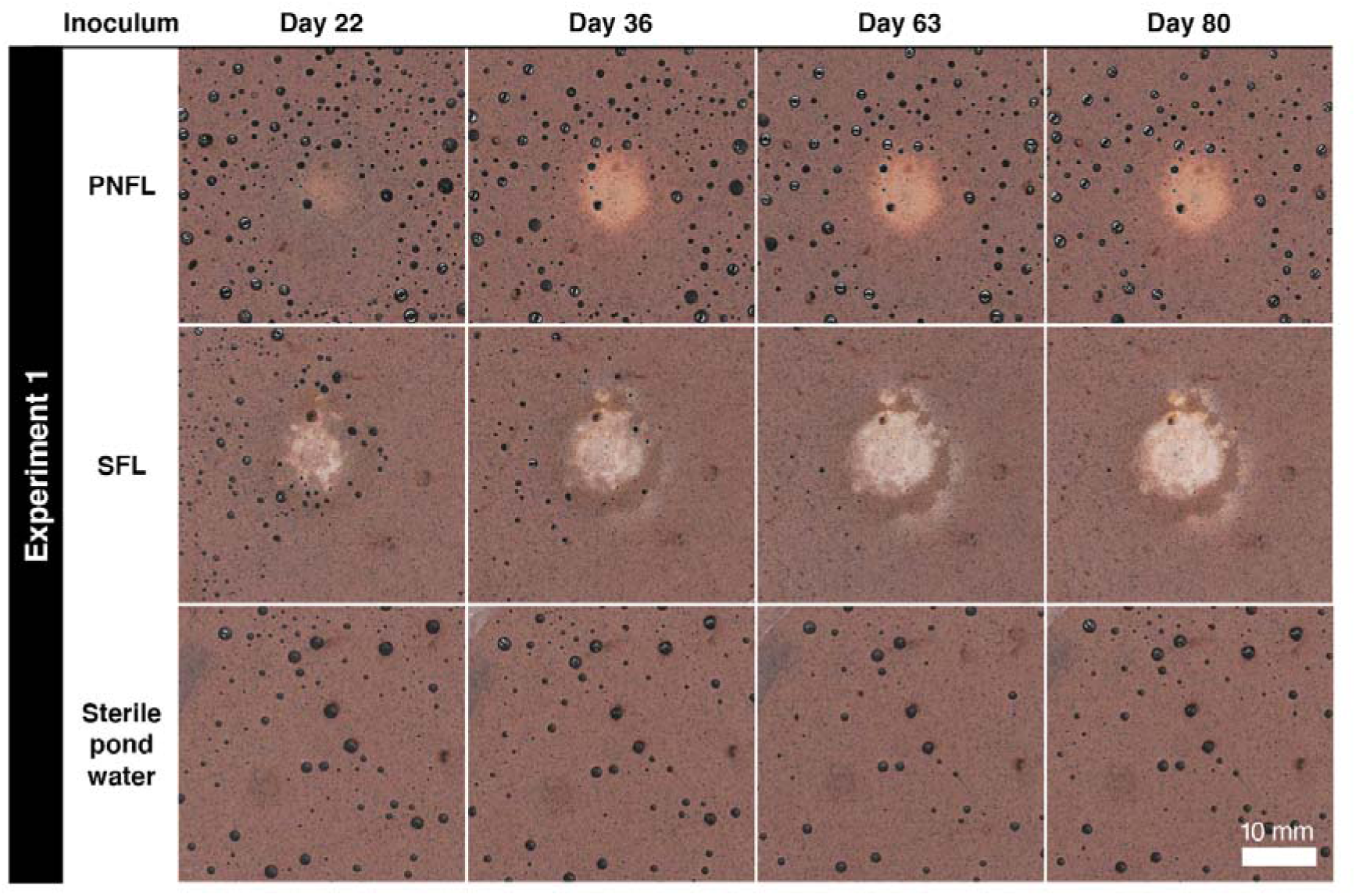
Time series of flat-bed scans showing central bleaching of the undersides of Petri dishes in Experiment. 1. Numbers of days refer to time since dishes were inoculated. Fields of view are aligned across time points. Dark circles are air bubbles. Spots developed directly beneath inoculated cotton wool in the dishes inoculated from Winogradsky columns PNFL (pond sediment, no ferrihydrite, light) and SFL (soil, ferrihydrite, light). The PNFL spot has less distinct margins than the SFL spot, which by day 36 had developed a secondary bleached halo at a remove of several mm from the original spot. Both PNFL and SFL spots showed some localized orange colouring to the margins.

**Table 1:** Occurrence of bleaching on the undersides of dishes. Numbers of days refer to time since Winogradsky columns were established. Note that bleaching occurred in all three dishes inoculated from the black layer in column SFL and did not occur when the inoculum from the same layer was filter-sterilized. The bleaching initially observed in the sample inoculated from column PNFL could not be replicated.

|  |  |
| --- | --- |
| **Experiment 1 (Day 57)** |  |
| PFL — unsterilized | No bleaching observed |
| PNFL — unsterilized | **Bleached spot(s) observed** |
| PFD — unsterilized | No bleaching observed |
| SFL — unsterilized | **Bleached spot(s) observed** |
| SNFL — unsterilized | No bleaching observed |
| SFD — unsterilized | No bleaching observed |
| Sterile pondwater | No bleaching observed |
| **Experiment 2 (Day 114)** |  |
| SFL — unsterilized | **Bleached spot(s) observed** |
| SFL — filter sterilized | No bleaching observed |
| PNFL — unsterilized | No bleaching observed |
| PNFL — filter sterilized | No bleaching observed |
| **Experiment 3 (Day 137)** |  |
| SFL — unsterilized | **Bleached spot(s) observed** |
| PNFL — unsterilized | No bleaching observed |
| **Experiment with *G. bemidjiensis*** |  |
| Unsterilized inoculum from liquid culture | No bleaching observed |

In SFL, the central spot was ∼8 mm across when first noticed and irregularly shaped with orange peripheral patches. By day 36 it had expanded to a ∼10 mm circle with well defined margins and a secondary bleached halo at a remove of several mm (**Figure 2**). By day 36, additional rust-orange material formed at the sediment-water interface along the margins of the same dish (**Supplementary Figure 2**); corresponding bleaching of the underside occurred in discrete ∼1-mm spots as well as larger blemishes (**Figure 3a**). Optical imaging **(Figure 3b)** and EDX mapping **(Figure 3c)** (after resin embedment) confirmed that the bleached zone was depleted in fine ferric particles compared to the matrix. No other elemental differences were apparent (e.g., in phosphorus or sulfur; **Supplementary Figure 3**).

**FIgure 3:**
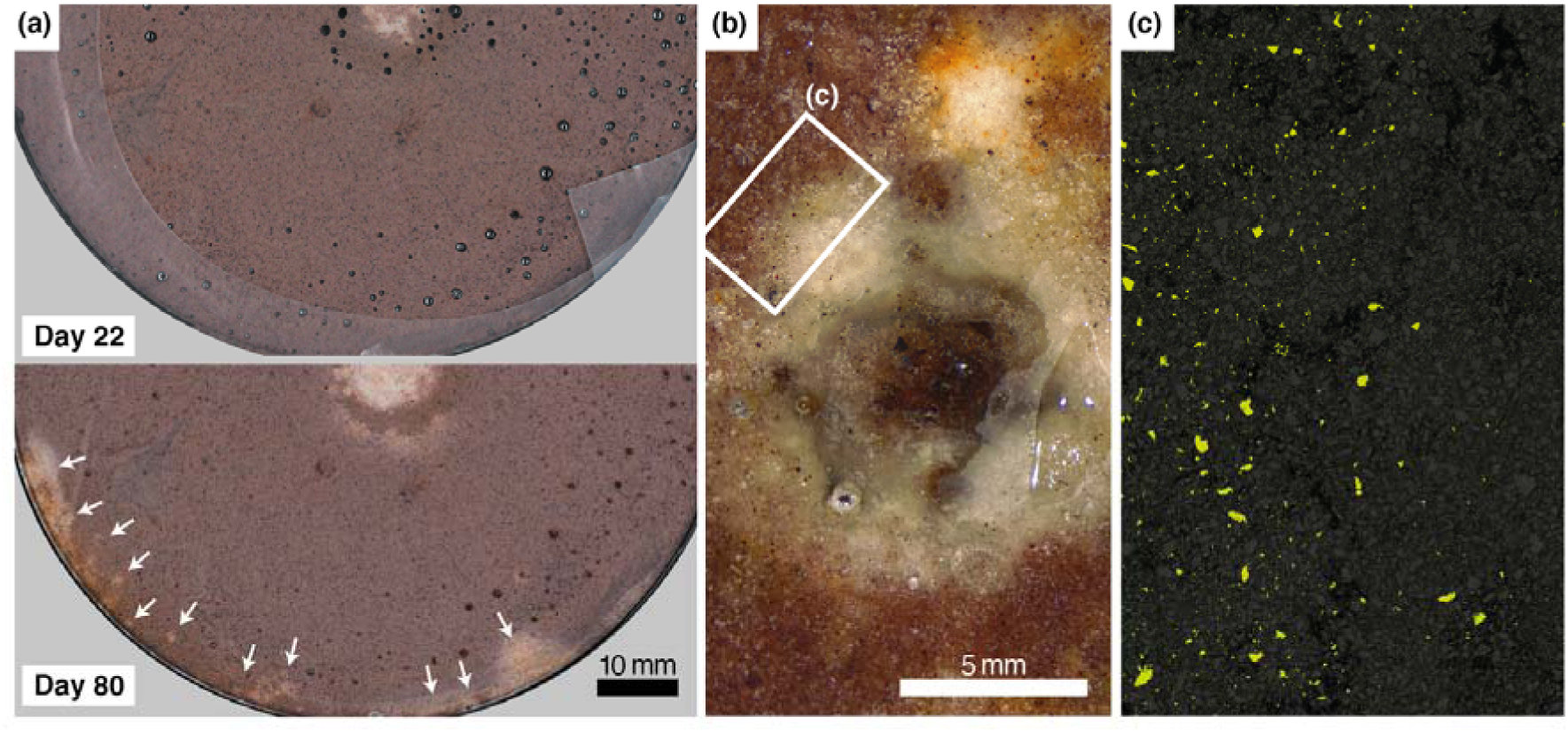
Additional observations from Experiment 1, dish SFL. **(a)** Wider field of view than Figure 2 showing development of new generation of bleached spots at dish margins by day 80 (white arrows), including discrete, mm-scale spots. Note associated orange-brown stain. **(b)** Photograph of initial spot from the same dish after resin embedment, grinding and polishing. During grinding, dark material inherited from the Winogradsky inoculum (not produced in the experiment) became visible. Bright flecks are reflections from air bubbles. Note orange-brown stain to upper right. **(c).** Overlaid EDX element maps from area highlighted in (b) of iron (bright green) and silicon (dark grey); remaining black areas indicate porosity filled by resin. Note decrease in iron abundance from visually red matrix into bleached zone.

In Experiments 2 and 3, bleaching occurred only in dishes inoculated from column SFL (**Figure 4**). Experiment 2 produced a faint, fuzzy spot with a dark orange-brown rim. An accompanying filter-sterilized control did not produce bleaching. Experiment 3 produced multiple spots with fuzzy margins and some orange colouring. No bleaching occurred in the dish with *G. bemidjiensis*.

**Figure 4:**
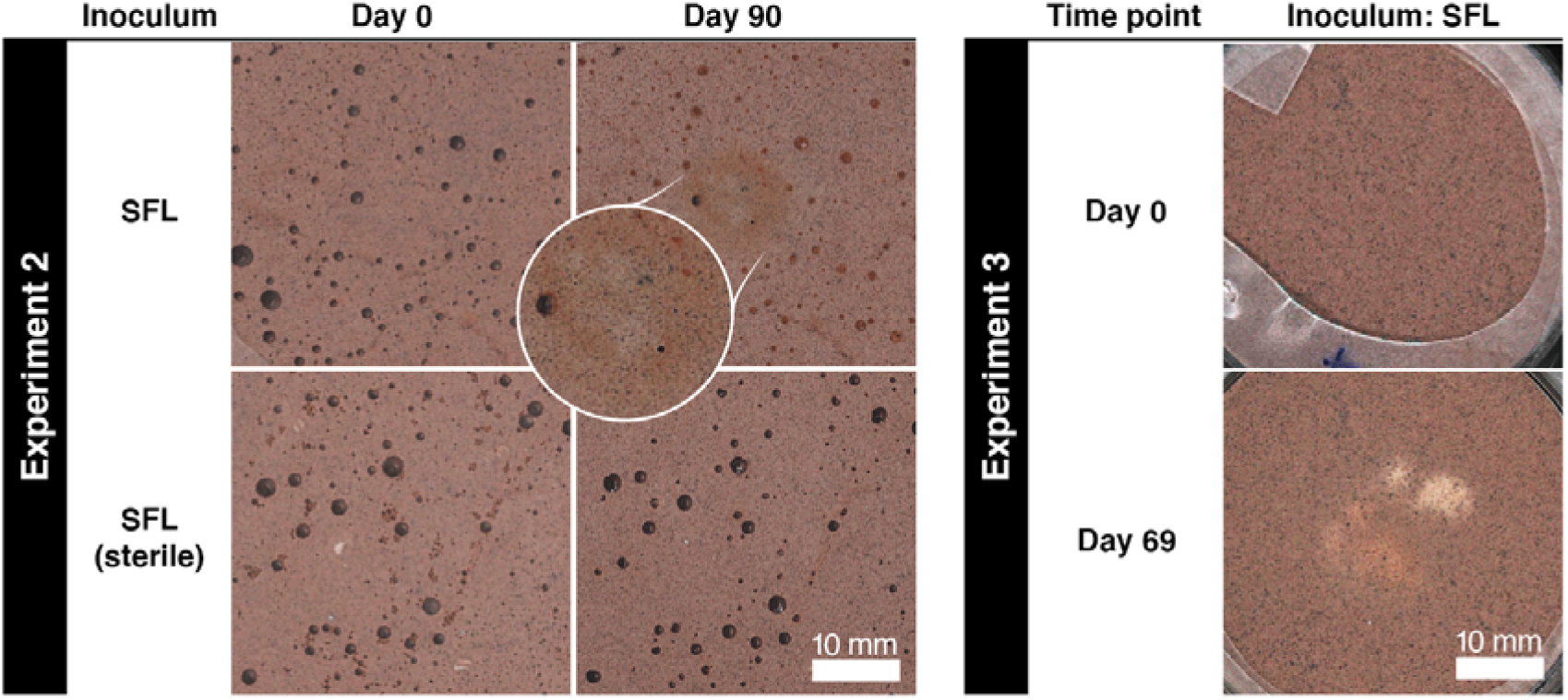
Flat-bed scans of Petri dishes showing bleaching in Experiments 2 and 3. Fields of view are aligned across time points. Experiment 2 produced a faint, fuzzy spot with a dark orange-brown rim. No bleaching occurred in the filter-sterilized control. Experiment 3 produced multiple spots with fuzzy margins and some orange colouring (40 mm Petri dishes were used for this experiment).

### Identification of iron-reducing bacteria

Liquid from Experiment 1 dish SFL was sampled from the sediment above the peripheral bleached zone (“side”) and from the wet cotton wool (“middle”) for 16S rRNA gene sequencing (see **Supplementary Information**). 488 amplicon sequence variants were identified across the two samples, indicating 136 distinct bacterial genera (**Figure 5**). Of those genera accounting for at least 1% of reads, genera typically characterised as Fe(III) reducers (*Paredesulfitobacterium* and *Desulfosporospinus*, both strictly anaerobic) jointly comprised 52.7% in “side” and 16.8% in “middle” (e.g., Li et al., 2021). Other genera containing at least some Fe(III)-reducing species comprised 30.2% in “side” and 36.6% in “middle”. Of these, the most abundant was *Cellulomonas* (a facultatively anaerobic cellulose degrader (Stackebrandt and Schumann, 2014) some of whose strains are known to perform dissimilatory Fe(III) reduction (Sani et al., 2002)). Genera not known to contain any Fe(III)-reducing species comprised 10.3% in “side” and 36.1% in “middle”. Genera accounting for <1% of reads comprised 6.8% of read in “side” and 10.5% in “middle”.

**Figure 5:**
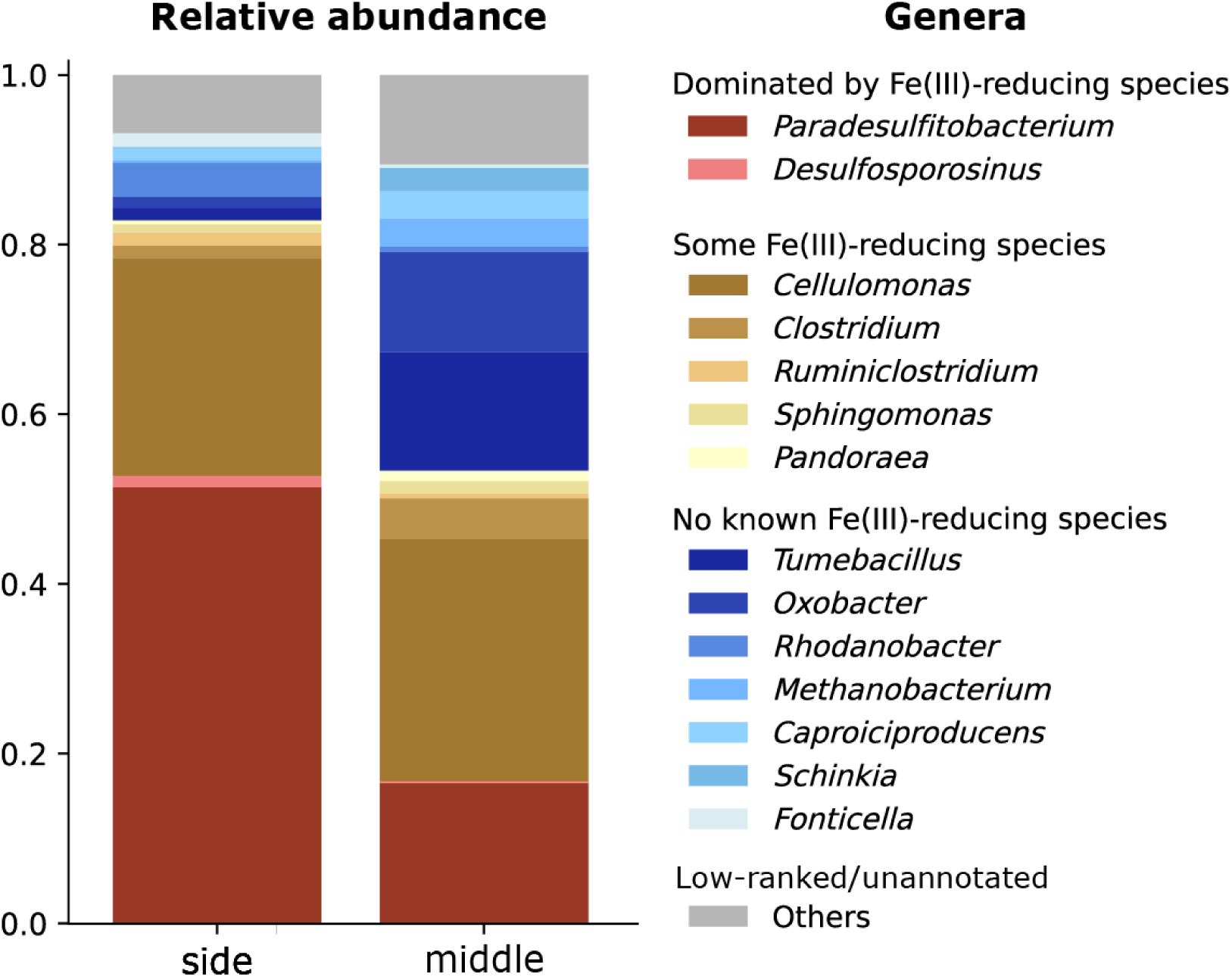
Genera occurring above reduction spots, and association with Fe(III)-reduction. Samples were taken from the sediment above the peripheral bleached zone (“side”) and from the cotton wool (“middle”) in dish SFL from Experiment 1. Genera of <1% abundance across both samples have been grouped in the category “Others”.

## Discussion

Our observations confirm that bleaching occurred via the reductive dissolution and diffusive loss of ferrihydrite. Orange-brown blemishes near the bleached zones and orange-brown crusts appearing on dish upper surfaces (**Supplementary Figure 2**) probably indicate re-oxidation of iron derived from the ferrihydrite. Vixseboxse et al. (2024) produced comparable orange halos around decaying organic matter in similar experiments. No other mineralization was observed.

Sulfides and organic acids can abiotically reduce and dissolve Fe(III), especially in poorly or non-crystalline phases, in days (Jauregui et al., 1982; Miller et al., 1986; Torres et al., 1990; Ionescu et al., 2015), (e.g., ferrihydrite; Miller et al., 1986). Organic acids adsorbed onto Fe-oxide surfaces form inner-sphere metal chelates; reduction via ligand-to-metal electron transfer results in the dissolution of the iron oxide and removal of ligand-bound Fe(II) complexes from the Fe-oxide surface, driving continued dissolution. Other than microbial biosynthesis, our experiments contained at least two sources of sulfides and organic acids, namely the 1 mL of material transferred from the Winogradsky columns and the pondwater in which all experiments were immersed. Nevertheless, we infer that bleaching observed in our experiments was microbially mediated, for five reasons:

(1) Of the four dishes inoculated from the same layer in column SFL, spots appeared in three, and failed to appear only in the one dish whose inoculum was filter-sterilized, a process that would have removed cells but not organic acids.
(2) The sequencing data revealed abundant bacteria of iron-reducing genera in the sediment above the bleached areas in SFL dish from Experiment 1 (64 days after inoculation); such genera were especially dominant in the peripheral area.
(3) All samples used to inoculate all six Petri dishes were reducing, sulfidic, and rich in organic matter, being taken from the black zone of Winogradsky columns. Nevertheless, inocula from only two columns caused bleaching, which must therefore require a more specific (and temperamental) agent than organic matter alone, e.g., certain microbial strains.
(4) Bleaching apparently accelerated in Experiment 1, suggesting a lag period typical of microbial growth kinetics. Abiotic solubilization of iron would have occurred rapidly after the introduction of material to the dishes and slowly thereafter (e.g., Miller et al., 1986). By contrast, dish PNFL bleached faster between days 22 and 36 than between days zero and 22 (Figure 1). Visible bleaching of the margin of dish SFL did not begin until after day 22.
(5) The late marginal bleaching observed in dish SFL occurred at multiple discrete mm-scale points as if induced by immobile localized agents such as microbial colonies (**Figure 3a**). Abiotic organic acids would not have been spatially focused in this way.

Although limited in size and scope, this study represents a first step towards a more exhaustive exploration of the mechanisms that might explain the occurrence of reduction spots in red rocks on Earth and Mars. Our general experimental approach is facile and easily replicated and extended. The method could be improved by replacing granular ferrihydrite with a more cement-like phase precipitated directly onto the sand grains. More generally, future work should seek diagnostically useful differences between bacteriogenic and abiotic reduction spots and their associated morphological, geochemical, and mineralogical fingerprints.

## Conclusion

We have observed that iron-reducing microorganisms can produce discrete, rounded, mm–cm-scale bleached spots in ferruginous sediments, suggesting that those found commonly in red beds on Earth—and those recently observed by *Perseverance* on Mars—are indeed “potential biosignatures”. However, the demonstration that microbes can produce reduction spots does not imply that all reduction spots are biogenic. It is now essential to search the abiotic baseline for non-biological pathways to reduction spot formation on Earth and Mars.

## Acknowledgments

We thank Kevin Dodd for resin embedding samples, Nicola Cayzer for assistance with SEM-EDX, and Charles Cockell, Niall Rodgers, and Seán Jordan for advice, support and discussion.

## Notes

### Competing Interest Statement

The authors have declared no competing interest.

### Funding

- Natural Environment Research Council: NE/Z504385/1

## References

Brown ST, Basu A, Ding X, et al. Uranium Isotope Fractionation by Abiotic Reductive Precipitation. Proceedings of the National Academy of Sciences 2018;115(35):8688–8693; doi: 10.1073/pnas.1805234115.

Gillen C, Jeancolas C, McMahon S, et al. The Call for a New Definition of Biosignature. Astrobiology 2023;23(11):1228–1237; doi: 10.1089/ast.2023.0010.

Hofmann BA. Reduction Spheroids from Northern Switzerland: Mineralogy, Geochemistry and Genetic Models. Chemical Geology 1990;81(1–2):55–81; doi: 10.1016/0009-2541(90)90039-a.

Hurowitz JA, Tice MM, Allwood AC, et al. Redox-Driven Mineral and Organic Associations in Jezero Crater, Mars. Nature 2025;645(8080):332–340; doi: 10.1038/s41586-025-09413-0.

Ionescu D, Heim C, Polerecky L, et al. Biotic and Abiotic Oxidation and Reduction of Iron at Circumneutral PH Are Inseparable Processes under Natural Conditions. Geomicrobiology Journal 2015;32(3–4):221–230; doi: 10.1080/01490451.2014.887393.

Jauregui MA and Reisenauer HM. Dissolution of Oxides of Manganese and Iron by Root Exudate Components. Soil Science Society of America Journal 1982;46(2):314–317; doi: 10.2136/sssaj1982.03615995004600020020x.

Lalla C, Calvaruso R, Dick S, et al. Winogradsky Columns as a Strategy to Study Typically Rare Microbial Eukaryotes. European Journal of Protistology 2021;80:125807; doi: 10.1016/j.ejop.2021.125807.

Li Y, Zhuang L, Yang G, et al. Paradesulfitobacterium Ferrireducens Gen. Nov., Sp. Nov., a Fe(III)-Reducing Bacterium from Petroleum-Contaminated Soil and Reclassification of Desulfitobacterium Aromaticivorans as Paradesulfitobacterium Aromaticivorans Comb. Nov. International Journal of Systematic and Evolutionary Microbiology 2021;71(9); doi: 10.1099/ijsem.0.005025.

McMahon S, Hood AvS, Parnell J, et al. Reduction Spheroids Preserve a Uranium Isotope Record of the Ancient Deep Continental Biosphere. Nature Communications 2018;9(1); doi: 10.1038/s41467-018-06974-9.

Miller WP, Zelazny LW and Martens DC. Dissolution of Synthetic Crystalline and Noncrystalline Iron Oxides by Organic Acids. Geoderma 1986;37(1):1–13; doi: 10.1016/0016-7061(86)90039-x.

Rogan B, Gorrell T, Lemke M, et al. Exploring the Sulfur Nutrient Cycle Using the Winogradsky Column. The American Biology Teacher 2005;67(6):348–356; doi: 10.2307/4451860.

Sani R, Peyton B, Smith W, et al. Dissimilatory Reduction of Cr(VI), Fe(III), and U(VI) by Cellulomonas Isolates. Applied Microbiology and Biotechnology 2002;60(1–2):192–199; doi: 10.1007/s00253-002-1069-6.

Seeger CH and Grotzinger JP. Diagenesis of the Clay-Sulfate Stratigraphic Transition, Mount Sharp Group, Gale Crater, Mars. Journal of Geophysical Research: Planets 2024;129(12); doi: 10.1029/2024je008531.

Smith SJ, Campbell BJ, Boerio-Goates J, et al. Novel Synthesis and Structural Analysis of Ferrihydrite. Inorganic Chemistry 2012;51(11):6421–6424; doi: 10.1021/ic300937f.

Spinks SC, Parnell J, and Bowden SA. Reduction Spots in the Mesoproterozoic Age: Implications for Life in the Early Terrestrial Record. International Journal of Astrobiology. 2010;9(4):209–216. doi:10.1017/S1473550410000273.

Stackebrandt E and Schumann P. The Family Cellulomonadaceae. In: The Prokaryotes. (Rosenberg E, DeLong EF, Lory S, Stackebrandt E, and Thompson F. eds) Springer: Berlin, Heidelberg; 2014; pp. 163–184; doi: 10.1007/978-3-642-30138-4_223.

Stylo M, Neubert N, Wang Y, et al. Uranium Isotopes Fingerprint Biotic Reduction. Proceedings of the National Academy of Sciences 2015;112(18):5619–5624; doi: 10.1073/pnas.1421841112.

Torres R, Blesa MA and Matijević E. Interactions of Metal Hydrous Oxides with Chelating Agents: IX. Reductive Dissolution of Hermatite and Magnetite by Aminocarboxylic Acids. Journal of Colloid and Interface Science 1990;134(2):475–485; doi: 10.1016/0021-9797(90)90157-j.

Vargas M, Kashefi K, Blunt-Harris E, et al. Microbiological Evidence for Fe(III) Reduction on Early Earth. Nature 1998;395(6697):65–67; doi: 10.1038/25720.

Vixseboxse PB, McMahon S and Liu AG. Taphonomic Experiments Fixed and Conserved with Paraloid B72 Resin via Solvent Replacement. Lethaia 2024;57(1):1–11; doi: 10.18261/let.57.1.1.

