## Supplementary Information for "Experimental evidence for a microbial origin of reduction spots in red beds"

*Images of Winogradsky columns*

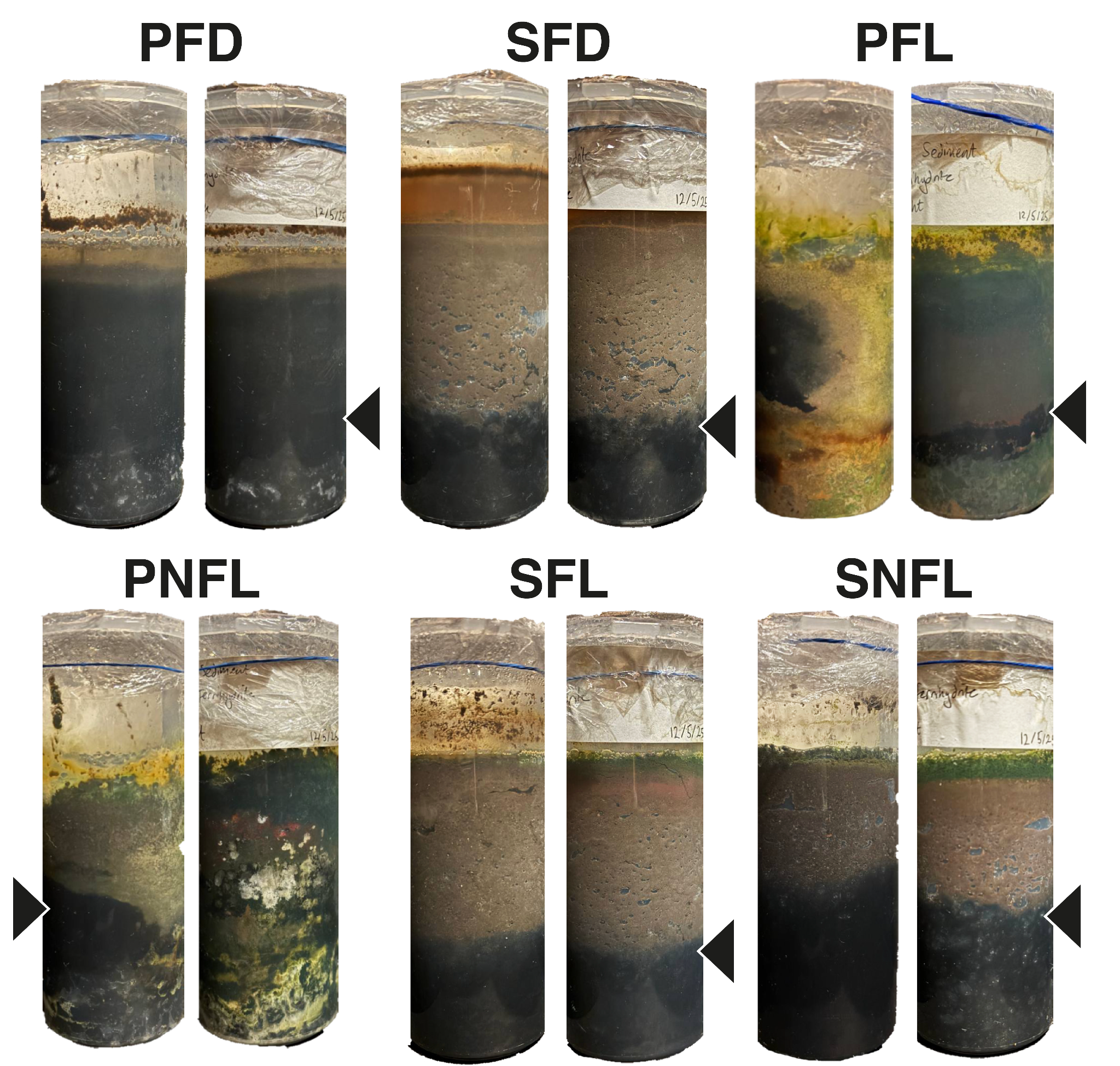

**Supplementary Figure 1: Front and back view of Winogradsky columns after 52 days.** Black sediment layers characteristic of microbial sulfate reduction are indicated by black arrows.

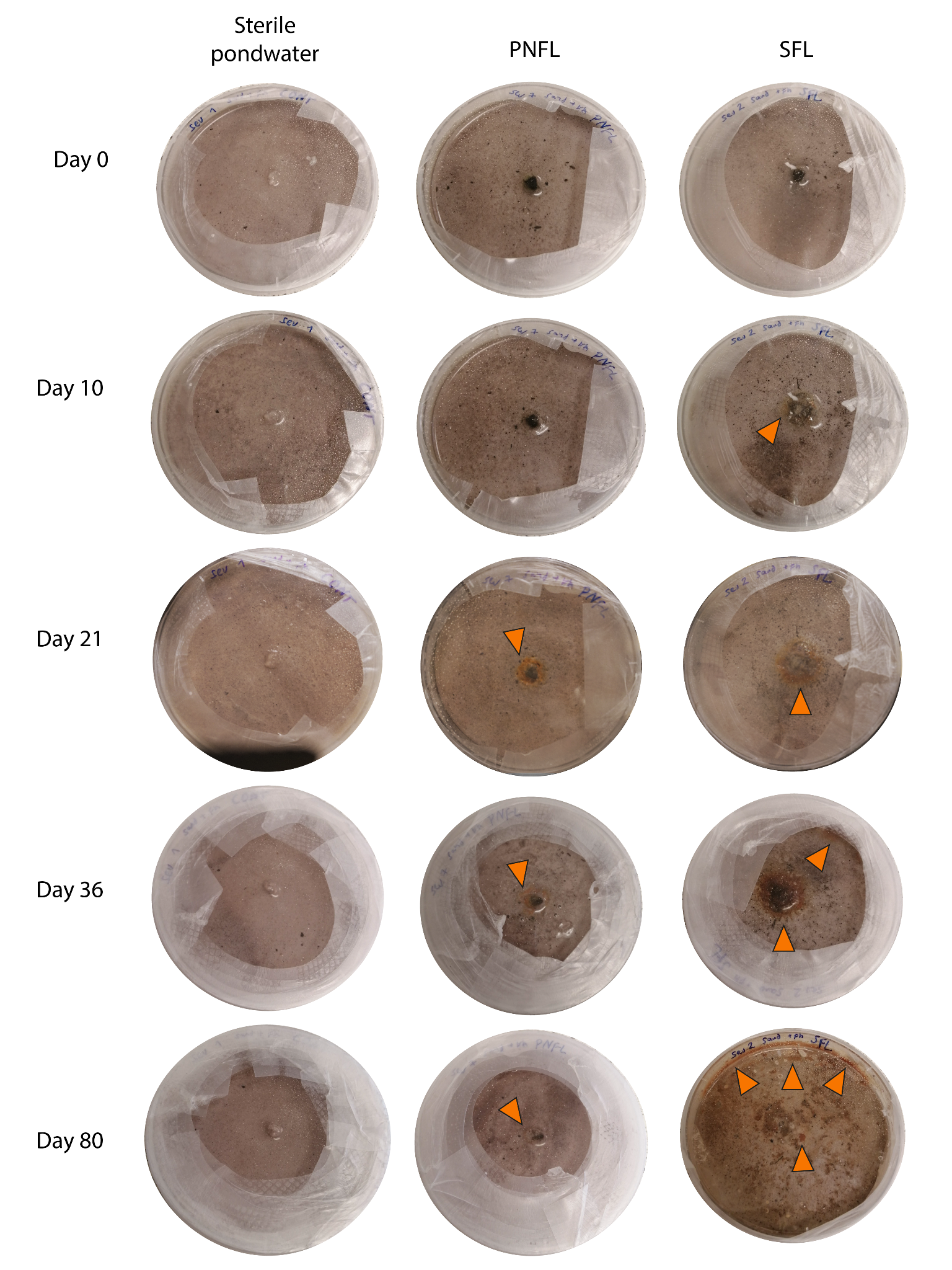

**Supplementary Figure 2** Upper surfaces of 90-mm Petri dishes photographed during Experiment 1. Orange arrows indicate orange patches, probably representing precipitation of re-oxidized iron. Other dishes behaved analogously to the sterile pondwater control.

*Resin impregnation methods*

To stabilize the reduction spot formed in the Petri dish inoculated from column SFL in Experiment 1, the sediment was first allowed to dry in the anaerobic chamber. The dish was then removed from the chamber and placed beneath a BVX-103 benchtop fume extraction system (Adhesive Dispensing ltd.) in the Thin Section and Sample Preparation Facility, School of GeoSciences, University of Edinburgh. A quantity of EpoThin 2 low viscosity epoxy resin was poured gently into the dry sediment and allowed to cure for twenty-four hours. The resin-impregnated solid disc was then freed from the Petri dish, cut down to size with a rock saw, and gently polished before further analysis. Some shrinkage of the central region of the reduction spot occurred during the drying or impregnation stage, creating voids that filled with resin. Polishing exposed a dark core region.

*Scanning electron microscopy methods and element maps*

The electron dispersive X-ray spectroscopy (EDX) data shown in Figure 3c were obtained with a Carl Zeiss SIGMA HD VP Field Emission scanning electron microscope fitted with an Oxford AZtec ED X-ray analysis system. A palladium coating was applied to the resin-embedded sample prior to electron microscopy. Element maps for S, Fe, Si and P are shown in Supplementary Figure 3.

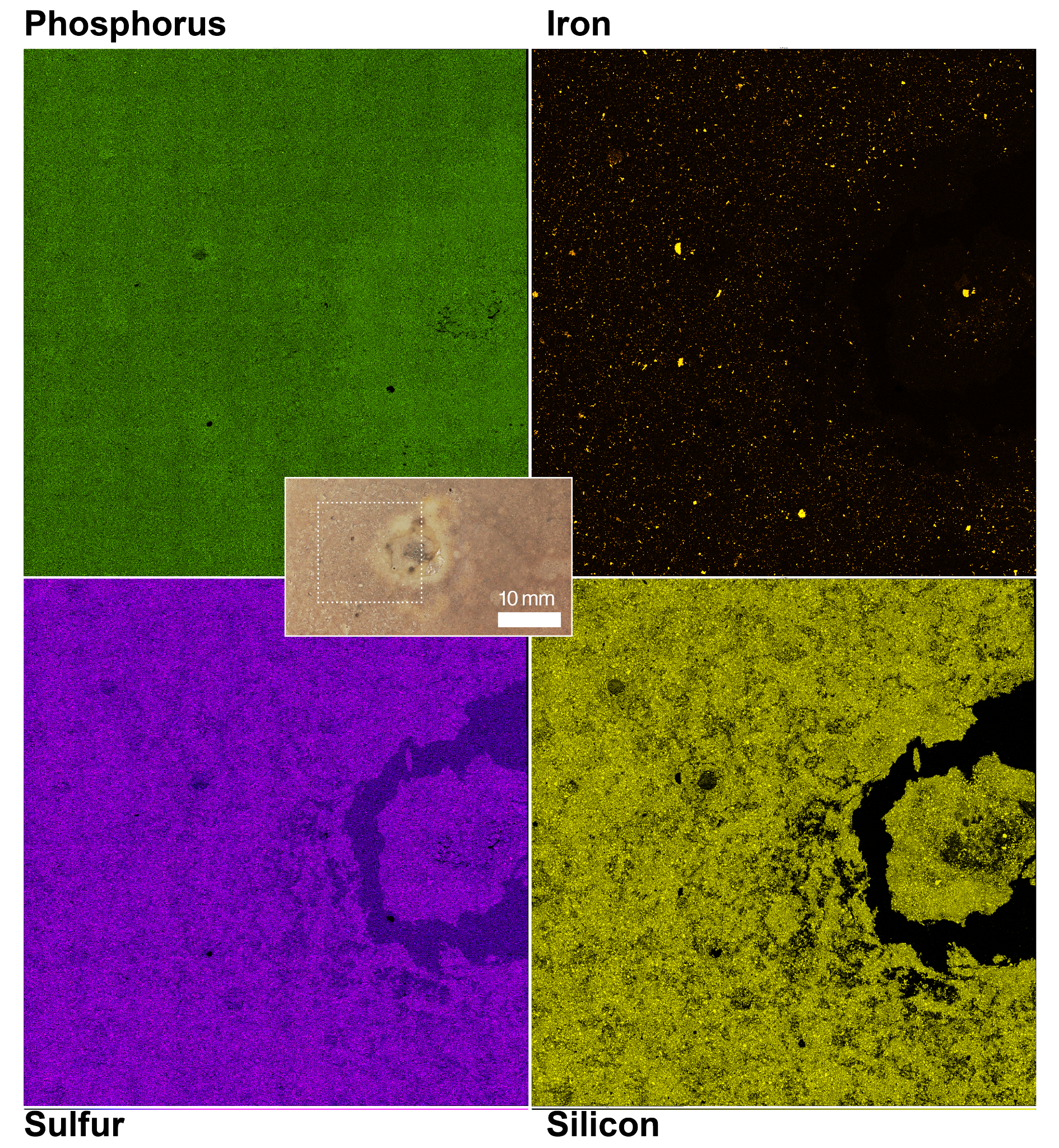

**Supplementary Figure 3: EDX element maps of iron, phosphorus, sulfur, and silicon (K-series).** The dark annular feature is a resin-filled void created during sample preparation. Note that resin appears to be slightly enriched in phosphorus compared to the sample. Inset shows mapped region of sample.

*DNA extraction and sequencing methods*

On day 64 after the initial Petri dish setup, two sediment samples were taken from SFL for DNA extraction. The first sample (“S” for “side”) was taken from the sediment at the dish margin where secondary bleaching had occurred. The second (“M” for “middle”) was taken from sediment on and around the cotton ball at the initial inoculation site. Samples were taken using disposable sterile inoculating loops inside of the anaerobic chamber.

DNA was extracted within 60 minutes of sampling using the DNeasy PowerSoil Pro kit (Qiagen), developed for use with soil and sediment. The nucleic acid (A260/A280) ratio was measured for each sample using a NanoDrop Microvolume Spectrophotometer (Thermo Scientific), resulting in a ratio of 1.71 for sample “M” (DNA concentration of 3.0 ng/uL), and 1.52 for sample “S” (DNA concentration of 2.1 ng/uL).

The DNA samples were stored at –20 C prior to postage to Novogene (Novogene Co., Ltd., Cambridge Sequencing Centre, United Kingdom), which performed sequencing and bioinformatic analysis of the samples. The following methodological details are condensed from information provided by Novogene.

PCR amplification of 16S rRNA genes in the region 16SV34 was carried out using primer sequences CCTAYGGGRBGCASCAG (forward mix) and GGACTACNNGGGTATCTAAT (reverse mix). PCR products were quantified and qualified according to Novogene procedure and sequencing libraries were generated and indexed. Quantified libraries were sequenced using the Illumina NovaSeq 6000 platform before passing through Novogene’s bioinformatics analysis pipeline:

Paired-end reads allocated to samples were combined using the FLASH analysis tool to form tags (V1.2.11, <http://ccb.jhu.edu/software/FLASH/>) (Magoc et al., 2011). Raw tags were quality-filtered using fastp software (V0.23.1) (Bokulich et al., 2012) and any chimaeral sequences were removed using the vsearch package (V2.16.0, <https://github.com/torognes/vsearch>) (Edgar et al., 2011) after comparison to the Silva reference database (<https://www.arb-silva.de/>) (Quast et al., 2013).

Effective tags were denoised (DADA2 module, Callahan et al., 2016), obtaining initial amplicon sequence variants, which could then be annotated taxonomically using QIIME2 software (Bolyen et al., 2019). QIIME2 was used to construct phylogenetic relationships through multiple sequence alignment.

Across the two samples, 206439 (S) and 202795 (M) raw paired-end (PE) reads were generated by the Illumina NovaSeq 6000 sequencer platform. Of these, 189250 (S, 91.67% of initial PE) and 178296 (M, 87.91% of initial PE) effective tags were used for downstream analysis, having removed any low-quality, short, ambiguous, or artificial sequence reads; 94.29% (S) and 94.89% (M) of high-quality (probability of an incorrect base call < 0.1%) PE reads were successfully assembled.

*Supplementary References*

Bokulich NA, Subramanian S, Faith JJ, et al. Quality-Filtering Vastly Improves Diversity Estimates from Illumina Amplicon Sequencing. Nature Methods 2013;10(1):57–59; doi: 10.1038/nmeth.2276.

Bolyen E, Rideout JR, Dillon MR, et al. Reproducible, Interactive, Scalable and Extensible Microbiome Data Science Using QIIME 2. Nature Biotechnology 2019;37(8):852–857; doi: 10.1038/s41587-019-0209-9.

Callahan BJ, McMurdie P, Rosen MJ, et al. DADA2: High-Resolution Sample Inference from Illumina Amplicon Data. Nature Methods 2016;13(7):581–583; doi: 10.1038/nmeth.3869.

Edgar RC, Haas BJ, Clemente JC, et al. UCHIME Improves Sensitivity and Speed of Chimera Detection. Bioinformatics 2011;27(16):2194–2200; doi: 10.1093/bioinformatics/btr381.

Magoč T and Salzberg SL. FLASH: Fast Length Adjustment of Short Reads to Improve Genome Assemblies. Bioinformatics 2011;27(21):2957–2963; doi: 10.1093/bioinformatics/btr507.

Quast C, Pruesse E, Yilmaz P, et al. The SILVA Ribosomal RNA Gene Database Project: Improved Data Processing and Web-Based Tools. Nucleic Acids Research 2012;41(D1):D590–D596; doi: 10.1093/nar/gks1219.
